# Joint processing of long- and short-read sequencing data with deep learning improves variant calling

**DOI:** 10.1101/2024.11.26.625492

**Authors:** Gennaro Gambardella

**Author notes:** CORRESPONDENCE, Address: Via Campi Flegrei 34, 80078 Pozzuoli (NA), Italy.

## Abstract

Despite the complementary strengths of short- and long-read sequencing approaches, variant calling methods still rely on a single data type. In this study, we collected and harmonized Nanopore dataset of the 7 GIAB healthy individuals across three independent consortia. Then, by leveraging these harmonized Nanopore data, we explore the benefits of using a hybrid DeepVariant model to jointly process Illumina and Nanopore data for germline variant detection. We show that a shallow hybrid long-short sequencing approach can match or surpass germline variant detection accuracy of state-of-the-art single-technology methods, potentially reducing overall sequencing costs and with the advantage of also enabling detection of large germline structural variations. These findings offer promising potential for molecular diagnostics in clinical settings, particularly for rare genetic disease screenings.

## INTRODUCTION

Although individual rare diseases might seem insignificant, their cumulative effect is substantial. Current data suggests that between 3.5% and 5.9% of the global population corresponding to roughly 300 millions people worldwide affected by rare diseases (1). Genetic factors are believed to cause approximately 80% of these rare conditions. Despite their widespread occurrence, most patients grappling with a rare or undiagnosed disease are limited to symptomatic relief, with only a scant 5% having access to appropriate treatments. Accurate diagnosis is the cornerstone in the effective handling of rare diseases. It enhances disease management and opens avenues for potential therapeutic interventions, all while avoiding unnecessary treatments that could result in severe side effects.

Over the past decade, next-generation sequencing (NGS), has significantly enhanced our ability to identify the causative variant and understand the inheritance pattern of rare genetic conditions. Initiatives like Care for Rare (https://www.care4rare.ca), Deciphering Developmental Disorders (2), Rare and Undiagnosed Diseases Diagnostic Service (3) and the Undiagnosed Diseases Network (4) have indeed widely demonstrated the transformative impact of whole exome (WES) and whole genome sequencing (WGS). However, it is crucial to acknowledge that only a fraction of patients (25-35%) currently receive a definitive molecular diagnosis followed by actionable findings.

Difficulties in pinpointing the causative variant from WES/WGS data are often related to detection challenges where the causative variant is in a difficult-to-sequence region, or it is obscured due to biases in reference genomes and genomic datasets. To this end, long-read sequencing provides more accurate and complete information about the structure and function of the genome, especially for regions that are difficult to resolve with short-read sequencing, such as repetitive or low-complexity regions.

In sum, short-read NGS technologies excel at identifying small variants but struggle in complex regions. Conversely, third-generation sequencing approaches based on long-read sequencing offer superior coverage of these regions, they excel at detecting large structural variants but have a higher error rate (i.e., 5-15%) that can compromise small variant detection (5). Indeed, even if advancements like Oxford Nanopore Technology (ONT) have currently reduced long-read sequencing costs of the human genome at 30X depth to about 850$ per individual and its last R10.4.1 chemistry can achieve a modal accuracy of Q20 for native reads, its high error rate compared to PacBio, severely limit its use in clinical practice for identifying small variants that are crucial for diagnoses of genetic diseases.

Essentially, both, short-read and long-read sequencing offer complementary advantages. However, current variant calling methods typically rely on a single data type, with limited attempts to combine PacBio and Illumina data (6). Therefore, hybrid approaches that can synergies the advantages of both sequencing approaches, correcting biases and enhancing accuracy, efficiency, and cost-effectiveness are urgently needed, especially for rare genetic diseases. Deep Learning (ML) and Artificial Intelligence (AI) have already emerged as powerful tools in the context of data integration in several biological fields (7). ML and AI algorithms can potentially correct errors and biases inherent in each sequencing method by creating a synergistic approach with higher resolving power. These approaches could enhance the accuracy and efficiency of variant detection and pave the way for a more cost-effective and timely analysis of rare genetic diseases. In this context, Convolutional Neural Networks (CNNs) have already been proven effective for variant calling tasks due to their ability to handle complex sequential data. Indeed, one such CNN-based tool, DeepVariant (8), developed by Google AI, effectively translates the problem of identifying variants into an image classification task, achieving high performance using short- or long-read data alone.

Here, to fully leverage the potential of both short-read and long-read sequencing, we developed an harmonized pipeline to process over 100 terabytes of publicly available raw ONT and Illumina sequencing data from the GIAB (9), HPRC (10), and the ONTOD project (11). These data were then feed to DeepVariant to train a novel model for identifying variants from hybrid short-long Nanopore-Illumina sequencing data.

We applied this hybrid model to several whole-genome case studies and show it outperforms state-of-the-art variant caller which solely relies on one sequencing type. We show that our hybrid approach can improve the accuracy of small variant detection, including the ones located in challenging repetitive/low-complexity regions. We show that a hybrid shallow whole-genome sequencing strategy combining 15X ONT and 15-X Illumina coverage can suffice to achieve a high germline variant detection accuracy, offering a promising solution for integrated small and large variant detection in clinical settings at a lower overall cost compared to deep sequencing performed with a single technology.

## MAIN

### A harmonized Nanopore dataset of individuals for training and benchmarking of variant caller algorithms

Public consortia like the GIAB (9), the HPRC (10), and the ONTOD project (11) provide valuable resources for developing and testing variant calling methods. These projects offer high-quality Nanopore and Illumina sequencing data for a well-characterized cohort of seven healthy individuals (HG001-HG007), including two trio families (9). Notably, the GIAB project includes a high confidence set of “ground truth” small germline variants for each individual, established by merging results from over 10 different sequencing methodologies (12, 13). However, data from these projects are fragmented and processed with different pipelines, potentially introducing biases that hinder their use for training AI-based methods.

To address these limitations, we built a harmonized Nanopore dataset specifically tailored for benchmarking variant calling algorithms. We collected over 100 terabytes of raw sequencing data in FAST5/POD5 format from all seven individuals across the three consortia (Table 01 & Supplementary Table 01). Data for Nanopore R10.4.1 chemistry was processed through a unified, publicly available pipeline we developed (https://github.com/gambalab/honey_pipes) optimized for this specific data type. This pipeline handles the entire workflow from raw reads to aligned BAM files, ensuring consistency and avoiding biases (Supplementary Figure 01 and Methods). Notably, the pipeline prioritizes simplex and duplex reads for precise allele frequency estimation, excluding redundant duplex parents reads (Methods). For Nanopore R9.4.1 data, we converted FAST5 files to FASTQ using Guppy and aligned reads with minimap2 (Supplementary Figure 01).

**Table 01.**
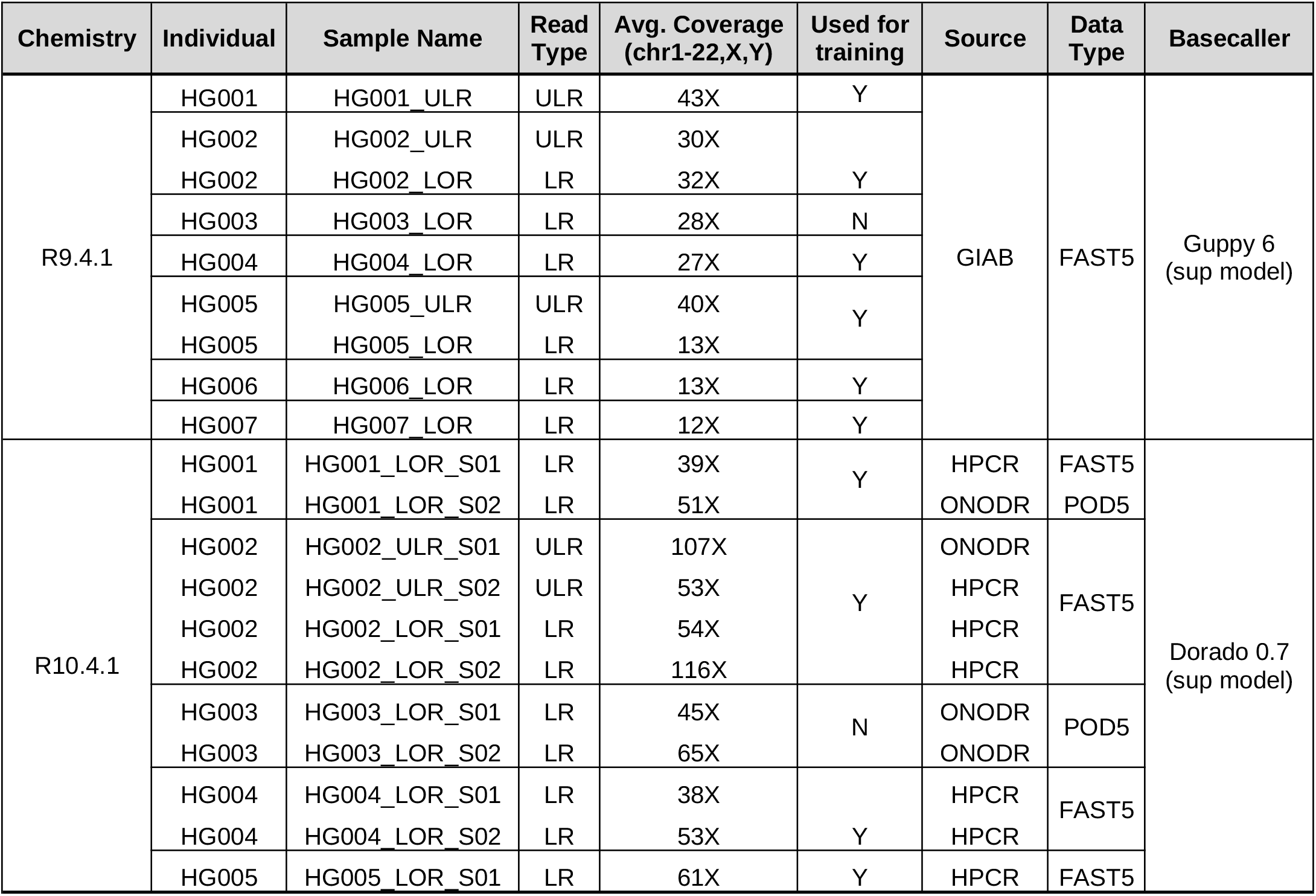
Collected Nanopore Samples.

The Nanopore datasets processed with these workflows, including both aligned and not aligned sequencing data for each individual, have been deposited on Sequence Read Archive (<u>see data availability section</u>) and are publicly available. These data offer a valuable resource for the scientific community to develop and test novel AI-driven variant-calling methods.

### Enhancing germline variant detection by jointly processing Nanopore-Illumina sequencing data

Leveraging the harmonized data we built, this study investigated whether a hybrid sequencing approach based on short-read Illumina data and long-read Nanopore data could improve small variant detection. To achieve this, we developed a custom Deepvariant model tailored to jointly process short- and long-read sequences from hybrid Nanopore-Illumina aligned data (available at https://github.com/gambalab/honey_deepvariant). Deepvariant (8) is a Convolutional Neural Network based variant caller have been developed by Google AI that translates the problem of identifying variants into an image classification task. We trained two novel Deepvariant models tailored to specific sequencing chemistries: (i) one adapted to recent Nanopore R10.4.1 chemistry and Illumina short-reads data and (ii) another adapted for old Nanopore R9.4.1 chemistry and Illumina short-reads data. We also employed data augmentation strategy using down-sampling (Methods) to increase the training dataset size that resulted in over 200 million training examples for each hybrid model (Table 01). Excluding HG003, the true germline variants from chromosomes 1-19 of the other collected individuals were used for training, while variants from chromosome 21 of each individual were used as validation data to evaluate model performance on unseen data during training stage.

Next, to evaluate our hybrid model’s performance, we applied them to a series of case studies where ground truth mutations are available from both GIAB consortia and ULTIMA genomics (14) and then compared our model’s accuracy to state-of-the-art variant callers applied on a single sequencing technology. Specifically, to explore the synergistic effect of combining short Illumina and long Nanopore reads, we simulated three whole-genome sequencing scenarios for HG003 individual. In the first scenario, HG003 was sequenced at a hybrid depth of 50X, combining 30X Nanopore and 20X Illumina reads. The second scenario used 30X Nanopore sequencing only, while the third used 20X Illumina sequencing only. As shown in Figure 1A-D and Supplementary Table 02, for single nucleotide variants (SNVs) detection, our hybrid model consistently outperforms leading Nanopore-only (Deepvariant (8), Clair3 (15), and Nanocaller (16)) and Illumina-only (GATK4 (17) and Strelka2 (18)) variant callers in terms of F1 score, true positive, false positive and false negative rates, regardless of used Nanopore chemistry and independently of the used ground-truth mutations dataset. While, for small INDELs detection (Figure 2A-D and Table 2), we observed a synergistic effect only when the latest Nanopore chemistry is used, especially when performances were evaluated on ULTIMA genomics ground truth mutation dataset. With previous Nanopore R9.4.1 chemistry, our hybrid model significantly outperformed Nanopore-only sequencing and performed comparably to Illumina-only variant callers, suggesting that in this case the model is not able to synergistically use both types of reads probably due to the highest error rate of R9.4.1 chemistry compared to R10.4.1 chemistry.

**Figure 1.**
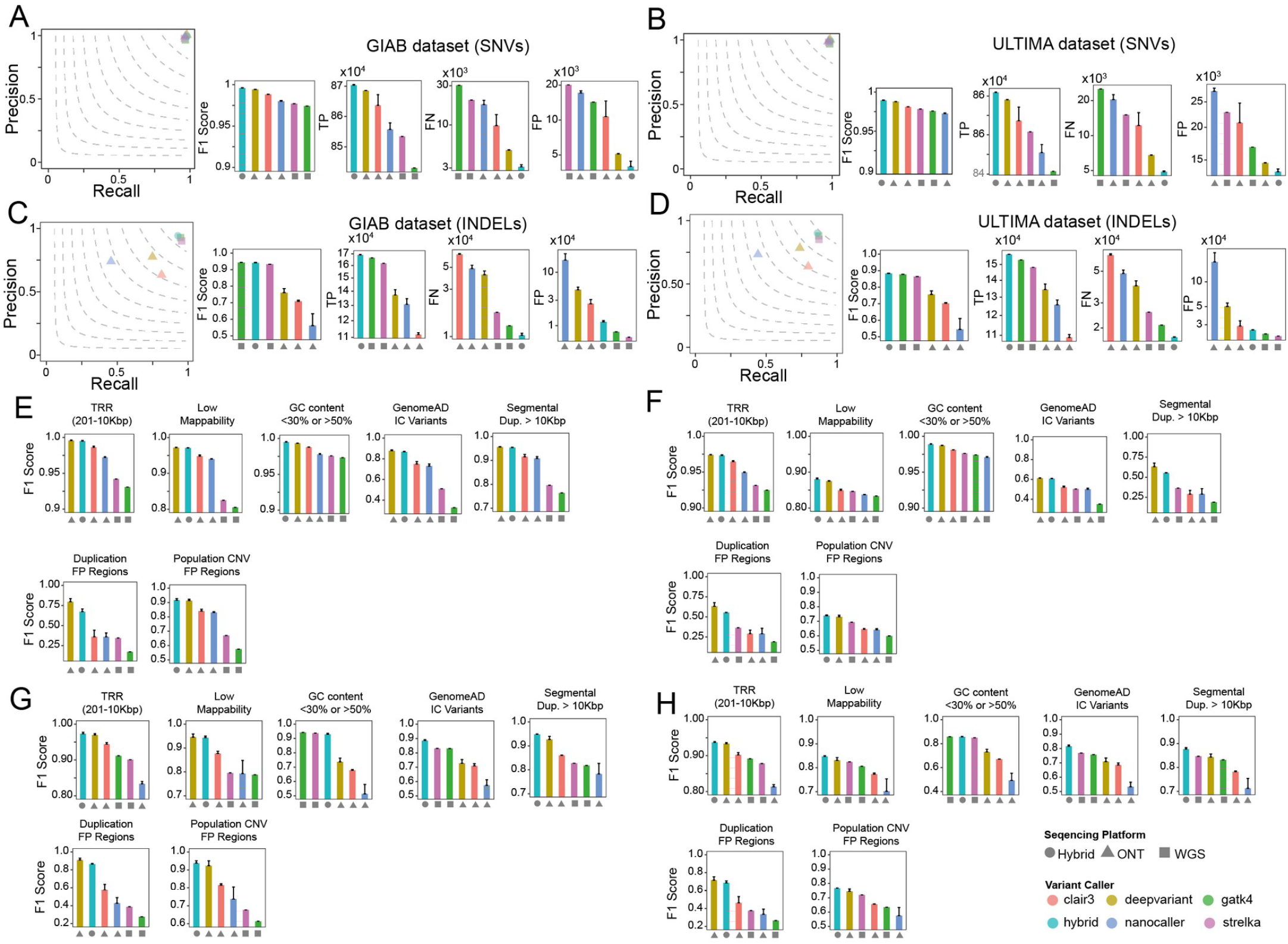
Combined Illumina and Nanopore sR10.4.1 chemistry Reads Enhance Germline Small Variant Detection. (**A**) Comparisons among Deepvariant Hybrid model and state of the art variant called relay on a single sequencing platform for detection of germline SNVs in individual HG003 using the GIAB ground mutations truth set. The Hybrid Model is tested on HG003, a mixture of 20x Illumina and 30x Nanopore reads. Single-platform callers (Nanopore only or Illumina only) use 30x Nanopore or 20x Illumina reads. TP = True Positives, FN = False Negatives, FP = False Positives. (**B**) Same as (A) but using ULTIMA as ground truth mutations set. (**C**) Same of (A) but for INDELs. (**D**) Same as (B) but for INDELs. (**E**) Comparisons among Deepvariant Hybrid model and state of the art variant called relay on a single sequencing platform for detection of germline SNVs in individual HG003 stratified by difficult genome regions. Performances are measured using GIAB ground truth mutations set (**F**) Same as (E) but using ULTIMA ground truth mutations set. (**G**) Same of (E) but for INDELs. (**H**) Same of (F) but for INDELs.

**Figure 2.**
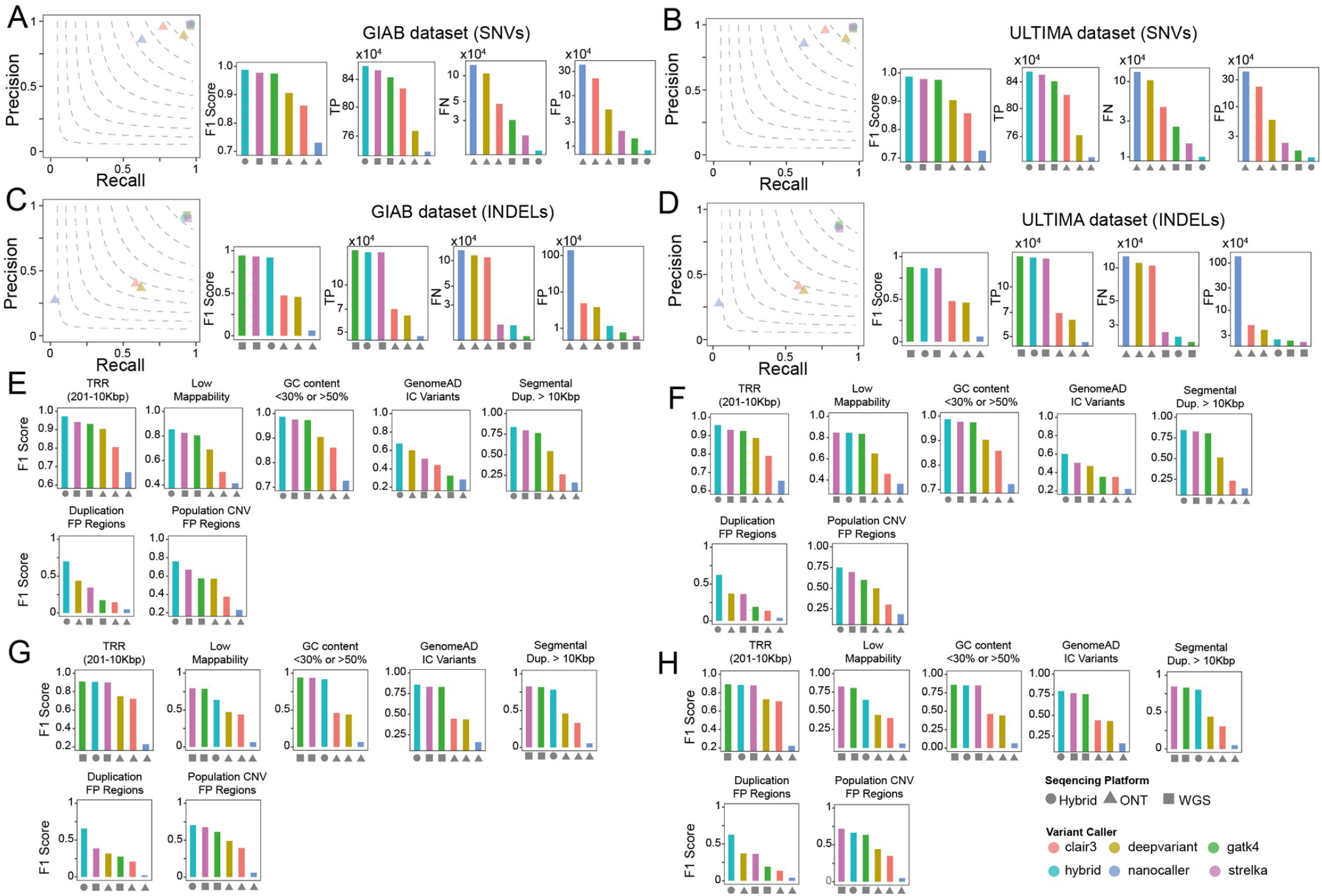
Combined Illumina and Nanopore R9.4.1 chemistry Reads Enhance Germline Small Variant Detection. **(A)**Comparisons among Deepvariant Hybrid model and state of the art variant called relay on a single sequencing platform for detection of germline SNVs in individual HG003 using the GIAB ground mutations truth set. The Hybrid Model is tested on HG003, a mixture of 20x Illumina and 30x Nanopore reads. Single-platform callers (Nanopore only or Illumina only) use 30x Nanopore or 20x Illumina reads. TP = True Positives, FN = False Negatives, FP = False Positives. Same as (A) but using ULTIMA as ground truth mutations set. (**C**) Same of (A) but for INDELs. (**D**) Same as (B) but for INDELs. (**E**) Comparisons among Deepvariant Hybrid model and state of the art variant called relay on a single sequencing platform for detection of germline SNVs in individual HG003 stratified by difficult genome regions. Performances are measured using GIAB ground truth mutations set (**F**) Same as (E) but using ULTIMA ground truth mutations set. (**G**) Same of (E) but for INDELs. (**H**) Same of (F) but for INDELs.

Next, we wondered if our hybrid model’s improved performance in detecting germline small variants was extended to challenging genomic regions. To this end, following GIAB guidelines (19), we stratified the genome of HG003 individual into seven challenging areas: Tandem Repeat Regions (TRR), Low Mappability Regions (LMR), GC enriched regions, regions enriched by variants with excess heterozygosity (i.e., which are defined by the InbreedingCoeff from the gnomAD database) (20), regions with long segmental duplications characterized by high sequence identity (21), and Collapsed Duplication FP regions and population CNV FP regions (related to collapsed duplication errors in GRCh38, recently identified and corrected by the T2T consortium (20)). As shown in Figure 1E-H and Figure 2E-H and Supplementary Table 02, we found that, regardless of the type of small variant or Nanopore chemistry used, our hybrid model either outperformed or almost matched the sequencing technology best suited for these challenging regions, showing the model’s ability to autonomously learn which sequencing technology to rely on for optimal variant detection in these regions.

Since the mutations of the HG002 individual (the son of HG003) are included in the training, approximately 50% of HG003’s variants are also used during the training of our method, as well as Clair3, DeepVariant, and Nanocaller. To correct this data leakage, all validation metrics computed for the HG003 individual presented above were assessed only after removing mutations shared with HG002 from the ground truth mutation set and predictions of each tool.

### A cost-effective whole-genome sequencing strategy using a shallow hybrid sequencing approach

Overall, these analyses demonstrated that combining long-read sequencing with R10.4.1 Nanopore chemistry with short-read Illumina sequencing can improve the accuracy of germline small variant detection, including the ones located in challenging repetitive or low-complexity regions of the genome. Therefore, next we asked whether we could identify the most cost-effective shallow hybrid sequencing protocol with an optimal trade-off between sequencing depth and variant detection accuracy. For this purpose, we used all available samples of the HG002 individual and generated several versions of its chromosomes 20 featuring different total hybrid coverage depths ranging from 10X to 35X (Figure 2A,B). In this case, we used chromosome 20 of HG002 individual due to the availability of the highest number of replicates for this individual. For these tests we used as ground truth the set of mutations the ones provided by GIAB. As shown in Figure3 A,B we found that after a total hybrid coverage depth of 25-30X the increase in F1 score of variant detection become modest (Supplementary Table 03), thus indicating that this depth could represents a good trade-off between sequencing depth and variant detection accuracy. However, only with chemistry R10.4.1 the SNVs detection accuracy was independent by the ratio of small and long reads used. In all the other scenarios the optimal trade-off was achieved within a range of 15X to 20X Illumina short reads and 10X to 15X Nanopore long reads (Figure 3A,B), indicating that while the two sequencing technologies can complement each other, the model has learned to prioritize short Illumina reads in these scenarios.

**Figure 3.**
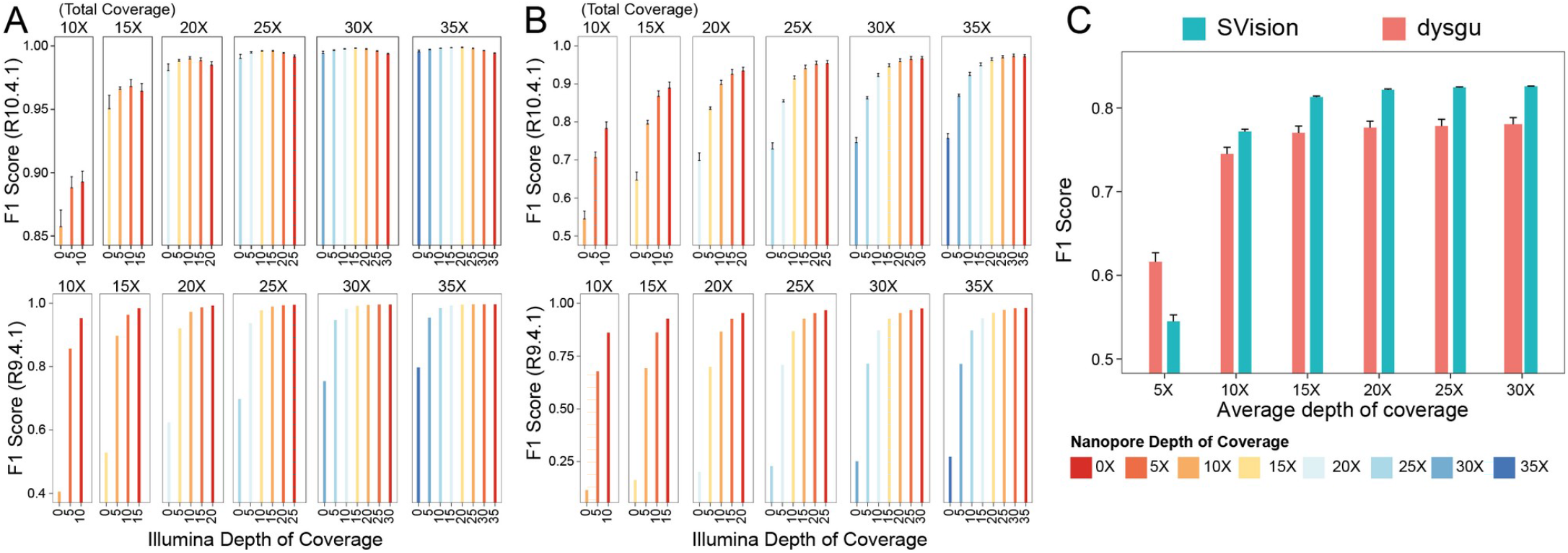
A shallow hybrid Nanopore-Illumina strategy for affordable whole genome sequencing. (**A**) How different ratio of Illumina and Nanopore reads affect performance of hybrid model for the detection of germline SNVs. Test were performed on chromosome 20 of HG002 individual. (**B**) Same as (E) but for germline INDELs. (**G**) Performance of dysgu and SVision tools in the identification of large SVs as a function of depth of coverage of whole genome of HG002 individual sequenced with Nanopore technology (chemistry R10.4.1).

Finally, to further demonstrate the detection accuracy of the shallow hybrid sequencing approach, we conducted a comprehensive comparison of DeepVariant models trained on hybrid sequencing data versus those trained on separate sequencing data. Specifically, for each of the seven GIAB individuals, we trained three distinct models: one using hybrid sequencing data, one using Illumina-only data, and one using Nanopore-only data. Each model was trained on chromosome 1, while variants from chromosome 21 were used as validation data to assess performance on unseen data during training. Model performance was then evaluated based on variant detection on chromosome 21 (Methods). All chromosomes were downsampled to a fixed total coverage of 30X (i.e., in the case of hybrid model we used 15X of short and 15X of long reads) and using Nanopore data from last chemistry R10.4. As shown in Supplementary Figure 02, our results confirm the synergistic advantage of combining short and long reads for SNVs detection. For INDELs detection, the hybrid model outperformed the Nanopore-only model while closely matching the performance of the Illumina-only model.

However, an advantage of long reads over short reads is also their intrinsic ability to detect large structural variations (SVs). Therefore, we wondered whether 10-15X of Nanopore long-read coverage would be also adequate for accurate SVs detection. To test this, we used all available samples of the HG002 individual for which ground truth SVs are available from GIAB and generated several downsampled versions of this individual with a depth of coverage ranging from 5X to 30X (see Methods). We then employed state-of-the-art AI-driven SV callers, such as Dysgu and SVision (22, 23), and assessed their performances as a function of Nanopore sequencing depth. As shown in Figure 2C, SVision consistently outperformed Dysgu, but for both methods the increase in F1 score beyond an average depth of 15X was modest (Supplementary Table 04), indicating that this depth of coverage is also likely sufficient for reliable structural variant identification from Nanopore long reads.

## CONCLUSIONS

In sum, this study demonstrates the value of harmonized datasets and hybrid sequencing approaches for improved variant detection. We created a comprehensive resource by collecting and processing raw Nanopore data from three different consortia (Methods). Additionally, we developed a set of state-of-the-art user-friendly pipelines for processing both Illumina and Nanopore data, accessible through a single Docker/Singularity container.

Leveraging this harmonized dataset, we trained custom DeepVariant models to jointly analyze short-read Illumina and long-read Nanopore data. Our method allows the generation of unified call sets rather than merging variant calls from different technologies. Our findings suggest that shallow hybrid sequencing (about 15X coverage for both platforms) achieves an F1 score of 0.9980 and 0.9454 in detecting SNVs and INDELs respectively. While structural variant with a depth of coverage of 15X Nanopore long-reads had a F1 score equal to 0.8129 with SVision (i.e., with 30X the F1 score was equal to 0.8259). Considering that a 30X Illumina whole genome sequencing costs approximately $1000, a hybrid sequencing approach with 30X coverage split equally between Illumina and Nanopore could potentially reduce the total cost to around $800, representing a cost reduction of approximately 20% while offering the advantage of covering both small and large germline structural variations. Additionally, although not explicitly demonstrated in this study, the Nanopore platform natively captures also promoter methylation status, further enhancing its utility. However, it is important to acknowledge that the proposed shallow hybrid sequencing approach would require access to two distinct sequencing instruments and necessitates separate library preparations, which also results in an increased requirement for the input genetic material.

While retraining existing single-technology methods like DeepVariant may have limitations, this study highlights the potential of combining short-read and long-read sequencing for improving variant detection particularly in complex regions of the genome. We present the first in-depth exploration of shallow hybrid sequencing for improved small variant detection. This approach holds promise for a more cost-effective and efficient solution in clinical settings like diagnosing genetic diseases. Our findings can be instrumental for the development of novel, native hybrid variant calling algorithms. These algorithms could accurately identify both small and large germline variants from shallow hybrid sequencing data, further reducing costs compared to deep single-technology sequencing.

## METHODS

### Nanopore R9.4.1 chemistry data collection and processing

We collected Nanopore sequencing data for all seven individuals (HG001-HG007) in FAST5 format from the GIAB consortium (links provided in Supplementary Table 01). We collected a total of nine samples, as HG002 and HG005 were sequenced with bot long and ultra-long reads approches (Table 01). Basecalling was performed using *Guppy v6* with the *super accurate* model, generating FASTQ files. Guppy was either run on our GPUs cluster equipped with 4 Nvidia A40 or Leonardo CINECA supercomputer equipped with custom Nvidia A100 GPUs. Alignment to the reference genome HG38 (primary assably from GENCODE) was achieved using *minimap2* with the *map-ont* preset alias.

### Nanopore R10.4.1 chemistry data collection and processing

Nanopore R10.4.1 FAST5/POD5 data for individuals HG001-HG005 were collected from HPRC and ONTOD consortia (links in Supplementary Table 01). We collected a total of 11 samples, as HG001, HG003 and HG004 sequenced with both long and ultra-long reads approaches while HG002 was sequenced in quadruplicate, two times using long reads and two times using ultra-long reads approach (Table 01). When starting with FAST5 format, data was first converted to POD5 with *pod5* phyton library. Subsequently, POD5 files were split by channels for optimized basecalling. Split POD5 files were basecalled with *Dorado v0.7.2* using the *super accurate* model in duplex mode, generating unaligned BAM files. We preferred BAM over FASTQ due to its inclusion of read type information (simplex, duplex or parent) crucial for accurate allele frequency estimation. Since FASTQ lacks this information, distinguishing read types is only possible with BAM input generated by the Dorado duplex basecalling command. To ensure precise allele frequency estimation, we strictly aligned only simplex and duplex reads, excluding redundant duplex parent reads that could distort variant calling. Read selection was performed with *samtools*. Dorado was run either on our GPUs cluster equipped with 4 Nvidia A40 or Leonardo CINECA supercomputer equipped with custom Nvidia A100 GPUs. Alignment to the HG38 reference genome (primary assably from GENCODE) was carried out using *minimap2* with the *lr:hqae* preset alias. Our pipelines for processing Nanopore data from FAST5/POD5 to aligned BAM files are publicly available on GitHub (https://github.com/gambalab/honey_pipes). These pipelines are encoded into the of three scripts *run_split_pod5.sh, run_dorado_duplex.sh* and *run_aln_long.sh*.

### Short Illumina reads data collection and processing

Illumina short-read FASTQ data for individuals HG001-HG007 was obtained from the National Institute of Standards and Technology (NIST) public FTP archive (ftp.ncbi.nlm.nih.gov). These data were used to generate hybrid short-long sequencing samples for training our DeepVariant hybrid model. Each flow cell offered an approximate 20X coverage depth. These FASTQ files were initially pre-processed using the *bbduk* tool (https://github.com/BioInfoTools/BBMap) to remove low-quality bases and adapter sequences. Subsequently, the pre-processed reads were aligned to the reference genome HG38 (primary assably from GENCODE) using our custom DRAGMAP aligner, available at https://github.com/gambalab/DRAGMAP. Finally, duplicate reads were filtered out using *samtools markdup*. The complete processing pipeline is encapsulated within the run_aln_short.sh script, accessible within our Docker/Singularity container at https://github.com/gambalab/honey_pipes.

### Training Hybrid Custom DeepVariant Models

We created hybrid sequencing datasets for training two DeepVariant models tailored to different Nanopore chemistries. For each individual, Nanopore sequencing data (aligned BAM files) was merged with Illumina data (aligned BAM files) using *samtools merge*. One dataset combined R9.4.1 chemistry Nanopore data with Illumina reads, while the other used R10.4.1 chemistry Nanopore data. To increase training data diversity, downsampled versions of each hybrid sample were created using *sambamba*, containing 50% of both short and long reads. We followed the official DeepVariant training guidelines (https://github.com/google/deepvariant/blob/r1.6.1/docs/deepvariant-training-case-study.md) and utilized chromosomes 1-19 from all hybrid samples for training. Chromosome 21 was reserved for validation, mirroring the approach of the original DeepVariant manuscript (8). Examples useful for training and validation were generated using Genome in a Bottle small variant true variants of each individual (v4.2.1) but only using truth confident region of chromosome 1-19 and 21. Training stage was employed using default channels 1-6 (version 1.6.1) and was carried out on machines equipped with Nvidia RTX 4090 and RTX 4080ti GPUs. After 10 epochs (approximately 11 days), training was stopped, and the best performing checkpoint was saved. A batch size of 1024 and 512 were respectively used for R9.4.1 and R10.4.1 hybrid models. For the hybrid model R9.4.1 258,665,415 examples were used for training and 3,455,573 for validation of the model during training. While for the hybrid model R10.4.1 210,962,694 examples were used for training and 2,584,574 for monitoring perfomces of of the model during training. Our custom DeepVariant code, specifically designed for hybrid Oxford Nanopore technology analysis (named “honey_deepvariant”), is publicly available as a fork on GitHub (https://github.com/google/deepvariant).

### Small Variant Callers comparisons on HG003 idividual (Figure 01 and Figure 02)

For Nanopore R10.4.1 chemistry, samples were downsampled to 30X coverage before before to be processed with Deepvariant v1.6.1, clair3 or Nanocaller. Downsamplig was performed with *sambamba* tool. While samples from R9.4.1 chemistry were used as they were since already had an average depth of coverage of about 30X (Table 01). Since no DeepVariant model exists for R9.4.1 chemistry, we employed pepper-DeepVariant (24) as recommended by the DeepVariant authors (https://github.com/google/deepvariant) to evalute Deepvariant performances on chemistry R9.4.1. The two short-read variant callers (GATK4 and Strelka2) were evaluated on Illumina data downsampled to 20X coverage (obtained from the GIAB consortium). Downsamplig was performed with *sambamba* tool. GATK4 followed best practices suggested by GATK, including Base Quality Score Recalibration around know SNPS and INDELs from 1000 genome project, while Strelka2 used standard parameters. GATK4 variant filtering was performed using the hard filtering strategy as per GATK guidelines (https://gatk.broadinstitute.org/hc/en-us/articles/360035531112--How-to-Filter-variants-either-with-VQSR-or-by-hard-filtering). To test performances of hybrid model bam files of a hybrid avarege coverage of 50X were instead built merging 30X from Nanopore and 20X from Illumina. Bam files merging was performed using *samtools*. Finally, performance of all variant callers, including our custom hybrid variant caller, was measured using the hap.py tool (https://github.com/Illumina/hap.py) in accordance with GIAB guidelines (12, 13). We analysed only *pass* mutations within confident regions for HG003, utilizing GIAB v4.2.1 small variant benchmarks or ultima genomics variants from s3://ultimagen-feb-2024-giab/DeepVariant_vcfs. Stratification files were obtained from GIAB consortia ftp server. To eliminate bias from training on HG002 (son of HG003), which includes about 50% of HG003’s variants, we removed shared variants from HG003’s ground truth and predictions before evaluating performance of each tool. This corrected the leakage present in our method, Clair3, DeepVariant, and Nanocaller which all have been trained using mutations of individual HG002.

### Identification of best ratio of long and short reads to enable shallow hybrid sequencing (Figure 03)

The different version of chromosome 20 of individual HG002 for all chemistry R10.4, R9.04 and Illumina were built first downsampling the original bam file and then merging the obtained downsampled Illumina and Nanopore bam files. Sambamba tool was used for donwampling while samtool for merging Illumina and Nanopore bam files.

### DeepVariant model comparisons (Supplementary Figure 02)

For each of the seven GIAB individuals, three distinct DeepVariant models were trained: one using hybrid sequencing data, one using Illumina-only data, and one using Nanopore-only data. Each model was trained on chromosome 1, while variants from chromosome 21 were used as validation data to assess performance on unseen data during training. The models were trained with a maximum of 50 epochs and an early stopping condition of 10 epochs. Model performance was then evaluated on chromosome 21 of the same individual. For both training and testing, all chromosomes were downsampled to a fixed total coverage of 30× (i.e., for the hybrid model, we used 15× short reads and 15× long reads). The Nanopore data used in to train hybrid models were the ones from R10.4 chemistry. The ground truth mutation set used to assess model performance was the GIAB v4.2.1 small variant benchmarks.

### Large Variant Callers Evaluation

We evaluated the performance of two structural variant (SV) callers, dysgu and SVision, on Nanopore R10.4.1 chemistry data for individual HG002. The four collected samples were downsampled to various depths of coverage (5X to 30X in 5X increments) using the sambamba tool. DYGUS and Snvsion were then run on each downsampled dataset with standard parameters. GIAB guidelines were followed for performance evaluation using Truvari (25). The SV benchmarks v1.1 for HG38 of individual HG002, available from GIAB project (https://www.nist.gov/programs-projects/genome-bottle), served as the truth set for this analysis.

## Declarations

### Ethics approval and consent to participate

Not applicable.

### Consent for publication

Not applicable.

### Availability of data and materials

Harmonized unaligned and aligned Nanopore data have been deposited on SRA with accession code PRJNA1191200 for R10.4 chemistry and PRJNA1193572 for R9.4 chemistry. Pipelines to process both short and long read are available at https://github.com/gambalab/honey_pipes, while our custom hybrid deepvariant tool is available at https://github.com/gambalab/honey_deepvariant.

#### Acknowledgments

We express our gratitude to IT core for supporting with cluster infrastructure.

## Authors’ Contribution

GG performed all bioinformatics analyses, wrote the manuscript and conceived the original idea.

## Declaration of generative AI and AI-assisted technologies in the writing process

During the preparation of this work the author(s) used gemini from google in order to proofreading the manuscript. After using this tool/service, the author(s) reviewed and edited the content as needed and take(s) full responsibility for the content of the publication.

