## Supplementary Figures for "Joint processing of long- and short-read sequencing data with deep learning improves variant calling"

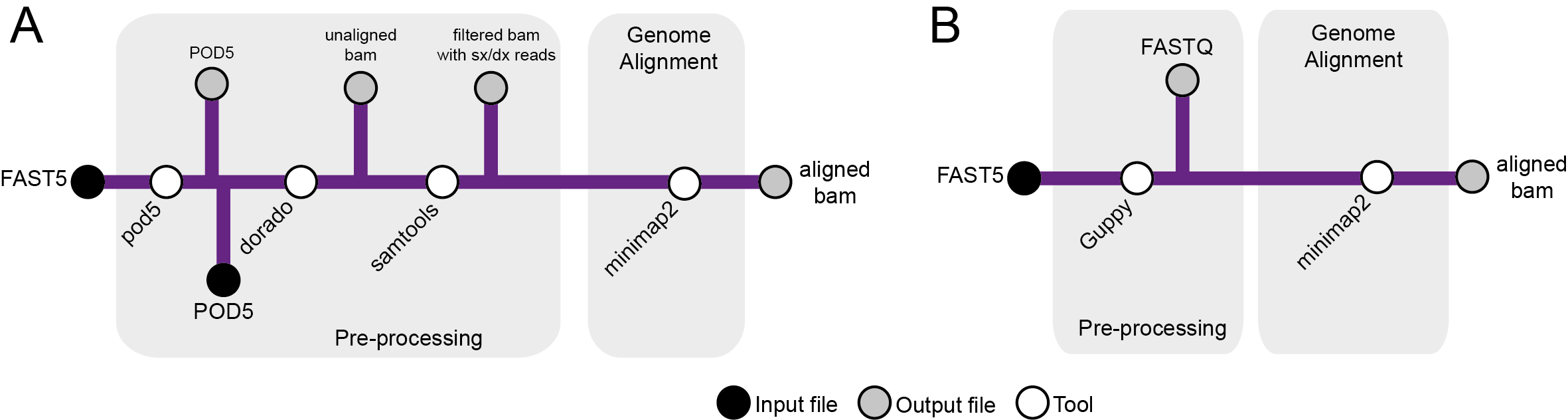

**Supplementary Figure 01 – Pipeline for harmonized Nanopore dataset.** (**A**) For Nanopore R10.4.1 data, FAST5 files are converted to POD5, split by channels, and then processed with Dorado for duplex basecalling. Samtools is used to filter out redundant duplex parent reads, retaining simplex and duplex reads. Minimap2 aligns these reads to a reference genome. (**B**) For Nanopore R9.4.1 data, Guppy converts FAST5 files to FASTQ, and minimap2 aligns the long reads.

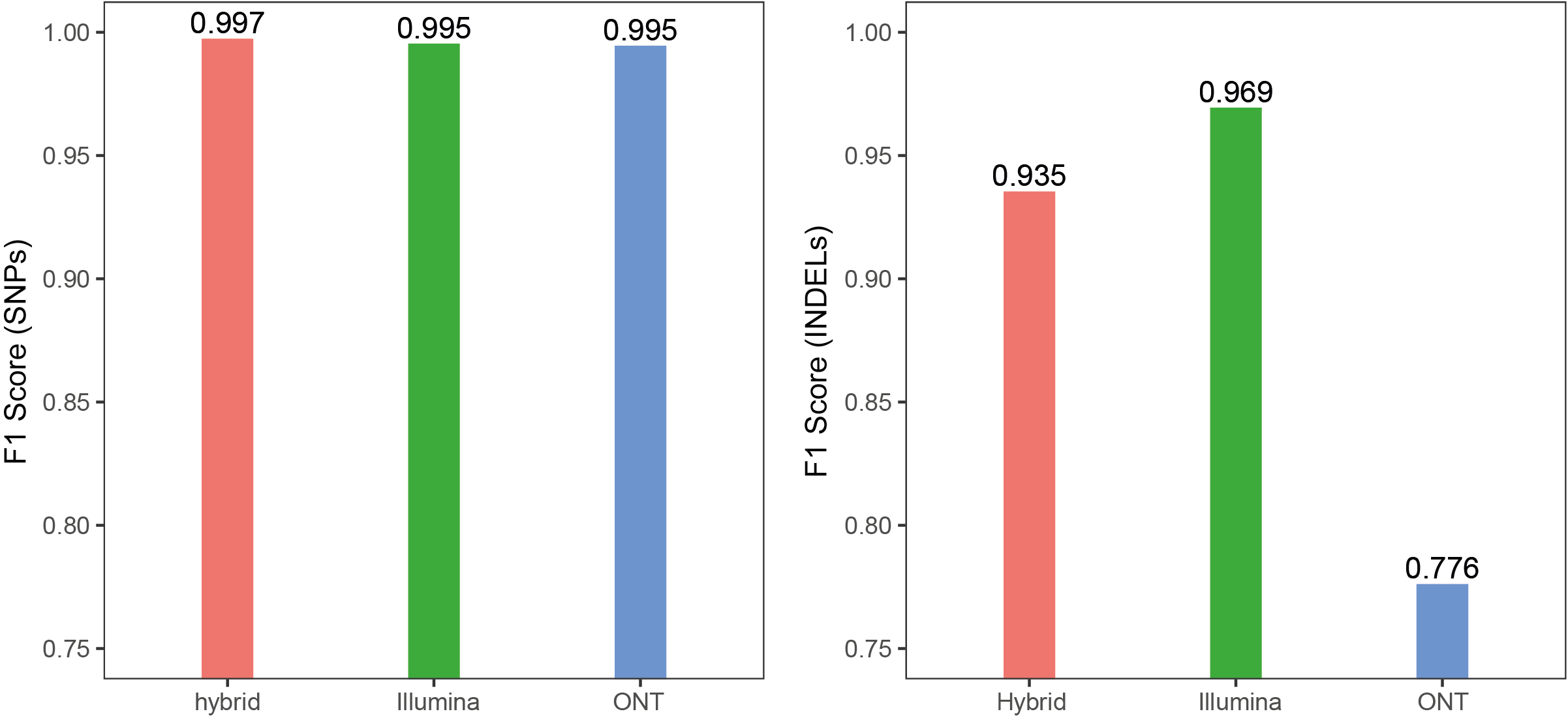

**Supplementary Figure 02 – DeepVariant model comparisons.** For each of the seven GIAB individuals, we trained three distinct DeepVariant models: one using hybrid sequencing data, one using Illumina-only data, and one using Nanopore-only data. Each model was trained on chromosome 1, while variants from chromosome 21 were used as validation data to assess performance on unseen data during training. Model performance was then evaluated based on variant detection on chromosome 21 of the same individual. All chromosomes were downsampled to a fixed total coverage of 30× (i.e., for the hybrid model, we used 15× short reads and 15× long reads) and analysed using Nanopore data from the latest R10.4 chemistry. (**A**) Average SNP detection performance of the three models on chromosome 21 across seven individuals. (**B**) Average INDEL detection performance of the three models on chromosome 21 across seven individuals. For both analyses, the ground truth mutation set was obtained from GIAB.
